# scSTAR2: a multiomics integration algorithm to reveal disease-specific cellular signatures by bridging single-cell resolution features and clinical metadata

**DOI:** 10.1101/2025.08.12.669818

**Authors:** Xin Zou, Jiawei Zou, Anzhuo Zhang, Fulan Deng, Yujun Liu, Xianbin Su, Henry H. Y. Tong, Lianjiang Tan, Wantao Chen, Jie Hao

**Affiliations:** Digital Diagnosis and Treatment Innovation Center for Cancer, Institute of Translational Medicine, Shanghai Jiao Tong University, Shanghai, China; Shanghai Institute of Biochemistry and Cell Biology, Center for Excellence in Molecular Cell Science, Chinese Academy of Sciences, University of Chinese Academy of Sciences, Shanghai, 200031, China; School of Materials Science and Engineering, Shanghai Institute of Technology, Shanghai 201418, China; Centre for Artificial Intelligence Driven Drug Discovery, Faculty of Applied Sciences, Macao Polytechnic University, Macao SAR, China; Department of Radiation Oncology, Fudan University Shanghai Cancer Center, Fudan University, Shanghai, China; Key Laboratory of Systems Biomedicine (Ministry of Education), Shanghai Center for Systems Biomedicine, Shanghai JiaoTong University, Shanghai, China; Ninth People’s Hospital, Shanghai Key Laboratory of Stomatology & Shanghai Research Institute of Stomatology, National Clinical Research Center of Stomatology, Shanghai Jiao Tong University School of Medicine, Shanghai, 200011, China; Shanghai Chenshan Plant Science Research Center, CAS Center for Excellence in Molecular Plant Sciences 201602, China

**Keywords:** single cell, ST, multi-omics integration, clinical meta data

## Abstract

Single-cell techniques, pivotal in characterizing intricate cell and spatial structures within tissues, face challenges in contextualizing clinical phenotypes. Currently, most investigations of the clinical phenotype related cell/spatial heterogeneities relied on the phenotype information from single-cell data. However, a wealth of underutilized clinical metadata exists within conventional bulk sequencing data. Current methods for correlating clinical information with individual cells or spatial regions remain limited and lack robustness. Here we present scSTAR2, a novel algorithm that transcends existing limitations by reconstructively integrating multiomics data to identify cells associated with clinical phenotypes. Unlike existing methods, scSTAR2 reconstructs single-cell data guided by specific phenotypes, significantly reducing interference from irrelevant noise. By employing scSTAR2 to integrate scRNA-seq, scATAC-seq, and spatial transcriptomics and bulk RNA-seq, we not only confirmed a new heat-shock Treg subtype in tumors but also more sensitively identified TLS (tertiary lymphoid structure) areas than traditional methods. In conclusion, scSTAR2 has proven to significantly enhance single-cell data interpretation across diverse clinical scenarios.

## Introduction

Single-cell technologies, including spatial transcriptomics, have revolutionized our understanding of cellular heterogeneity, offering key insights into cell types, states, and spatial organization. Unsupervised clustering is a common approach for identifying cellular subpopulations, analyzing functions, and inferring cell-cell interactions [1, 2]. Identifying clinically relevant subpopulations, such as those linked to tumor progression or therapy response, is pivotal for advancing personalized medicine [3].

Efforts to characterize phenotype-associated subpopulations using single-cell data often involve comparing cellular compositions across phenotypes [4-6]. For example, in our previous study [7], we identified immunotherapy-linked cell subtypes using GSVA to allocate differentially expressed (DE) genes between sensitive and resistant samples in specific clusters [8]. However, the high costs and stringent requirements of single-cell techniques limit their scalability in large-cohort studies, posing challenges for broader clinical applications.

Conversely, bulk RNA-seq datasets, such as TCGA [9] and GEO [10], represent decades of accumulated data and contain rich clinical phenotype information. Bulk RNA-seq has been used to associate clinical features like drug response [11-13], prognosis [14, 15], and relapse[16] with molecular classifications. Nonetheless, bulk methods lack cellular resolution, resulting in potentially obscured or compromised outcomes.

To bridge bulk and single-cell data, methods like Scissor [17], scDEAL [18], and scAB [19] integrate bulk RNA-seq with scRNA-seq to identify phenotype-associated subpopulations. However, their reliance on raw scRNA-seq data, often impacted by noise unrelated to the phenotypes of interest, compromises precision. Our previous work, scSTAR [22], addressed this issue by reconstructing single-cell profiles based on phenotype-specific dynamic patterns, demonstrating robustness to noise [23, 24]. Nonetheless, scSTAR’s reliance on phenotype-specific single-cell datasets limits its applicability in diverse clinical and biological contexts.

To overcome these limitations, we present scSTAR2, a novel algorithm that integrates single-cell and bulk RNA-seq data to extract phenotype-associated gene expression dynamics. scSTAR2 exploits abundant bulk RNA-seq data with clinically relevant phenotypes to improve single-cell data interpretation. By projecting scRNA-seq, spatial transcriptomics, or scATAC-seq data into a low-dimensional space informed by phenotype data, scSTAR2 filters out irrelevant noise while preserving phenotype-relevant components. The data is then reconstructed in this refined space, enabling robust phenotype-cell mapping. Unlike scSTAR, scSTAR2 eliminates the need for collecting single-cell data from samples with distinct phenotypes, significantly broadening its applicability.

Benchmarking revealed that scSTAR2 outperforms existing phenotype-mapping tools across both simulated and experimental datasets. It successfully disentangles clinically meaningful components from single-cell data under diverse phenotypic contexts, facilitating interpretation and highlighting the translational potential of single-cell technologies in clinical research.

Single-cell techniques (including spatial transcriptomics) have revolutionized our understanding of transcriptomic variations between cells, revealing critical insights into cell types, states, and spatial characteristics. To characterize cell heterogeneities and elucidate inter-cellular relationships, unsupervised clustering is usually performed to categorize cells into clusters, investigate the corresponding cellular functions and infer the cell-cell relationships [1, 2]. Identifying cell subpopulations linked to specific clinical phenotypes, including tumor prognosis and treatment response, is crucial for advancing personalized medicine. [3].

Efforts have been made to investigate the phenotype associated cell subpopulations using single-cell techniques, which all require the single-cell data acquired from the tissues associated with different phenotypes. One popular way is to interrogate the cellular composition changes between different phenotypes [4-6]. In our previous work [7], we identified the immunotherapy treatment associated cell subtypes by allocating the DE genes between sensitive and resistant samples on specific cell cluster using GSVA [8].

However, the high cost and stringent quality requirements of single-cell techniques limit their applicability in large cohort studies, restricting their potential in broader clinical research.

Conventional bulk RNA-seq database, accumulated over decades, holds a wealth of clinical phenotype information, presenting an untapped resource for molecular classification and clinical correlations, e.g., The Cancer Genome Atlas (TCGA) [9] and Gene Expression Omnibus (GEO) [10]. With bulk RNA-seq data, clinical phenotypes can be associated with molecular classification, e.g., drug sensitivity [11-13], prognosis [14, 15], relapse [16]. However, bulk technique-based studies lost abundant cell details and may be just compromised results.

To leverage the single-cell technique guided cancer molecular classification, computational techniques have been developed to transfer the bulk-level phenotype information to single-cell level very recently. Scissor [17] identifies cell subpopulations from single-cell data associated with a given phenotype by integrating phenotype associated bulk RNA-seq data with scRNA-seq data. This is achieved by constructing a regression model between the cell-sample correlation matrix and the sample phenotype. scDEAL [18] uses deep transfer learning model to predict the drug response patterns for each cell based on bulk RNA-seq and scRNA-seq data. scAB [19] integrates scRNA-seq data with labeled bulk data by knowledge- and graph-guided matrix factorization model, but is limited by prior-knowledge of phenotype associated pattern, which typically corresponds to a known biological process/signal associated with the phenotype of interest. In addition, PACSI [20] and DEGAS [21] were also developed to associate clinical phenotypes with single cells.

All of the aforementioned methods performed phenotype-cell mapping using the original scRNA-seq data. As it has been known, the variations contained in scRNA-seq are complicated, most of which are irrelevant to the phenotypes of interest. It is always a challenge to extract useful information from noisy data. In our pervious study, we developed scSTAR [22] to reconstruct single-cell data on the cell dynamic patterns extracted with the guidance of phenotype information from single cell data. scSTAR showed high noise robustness and was successfully applied in many clinical scenarios [23, 24], but the single-cell-data-phenotype dependency hampered its applications in more general cases.

Building on the foundation laid by scSTAR, we introduce scSTAR2, a novel algorithm that extracts phenotype-associated gene expression dynamics from various types of single-cell data, harnessing the synergy of single-cell data and bulk RNA-seq data integration. scSTAR2 makes full use of the widely accessible bulk RNA-seq data with clinically valuable phenotype information. With the guidance of phenotype information from independent data, scSTAR2 first projects the original single-cell data e.g., scRNA-seq, spatial transcriptomics and scATAC-seq, into a low dimension space, in which only the components related to the phenotype alterations are maintained. Then, the data is recovered from the low dimension space, which can filter out irrelevant noises. In the framework of scSTAR2, phenotype-cell mapping is performed on the reconstructed data. Compared to scSTAR, scSTAR2 no longer requires the single cell data collection from the samples of different phenotypes. Furthermore, scSTAR2 outperformed existing phenotype-cell mapping methods on both simulated and experimental data. Our studies demonstrate that scSTAR2 can dissect various clinical meaningful components hidden in a given single-cell dataset with the guidance of various phenotypes, which not only facilitates the single-cell data interpretation, but also sheds light on potential clinical applications of single-cell techniques.

## Results

### Design of scSTAR2

Building on our previous scSTAR [22] findings, we identified the need for a method less reliant on the phenotype information from single-cell data, leading to the development of scSTAR2. This new version capitalizes on the rich clinical data in bulk samples, extracting cellular dynamics correlated with phenotypes. Similar to scSTAR, scSTAR2 is engineered to extract cellular dynamics associated with studied phenotypes. However, it uniquely reconstructs single-cell data guided by clinical phenotype information from bulk RNA-seq data, enhancing its applicability to clinical scenarios. Therefore, the input of scSTAR2 consists of a query scRNA-seq expression matrix (or other single-cell resolution omics data), an independent phenotype-provider data, e.g. bulk expression matrix with categorizable phenotype information. Currently, scSTAR2 only supports the cases of binary phenotypes, which resonates with prevalent clinical scenarios like good/poor prognosis, drug responses, and metastatic events.

scSTAR2 operates in several key steps: 1) Extraction of common features between the query single-cell and the phenotype-provider data in a shared latent space using a partial least square (PLS) model, Then, reconstruct the query single-cell and phenotype-provider data using the shared features only, respectively; 2) Building a second PLS model (PLS2) to relate the phenotype-provider data to phenotypes. PLS2 is only sensitive to the gene expression variation components which associated with the changes of phenotypes; 3) Reconstructing query single-cell data to remove irrelevant gene expression variations. The reconstructed query single-cell data are termed as ‘pseudo-dynamic data’; 4) Determining phenotype-cell associations using an orthogonal partial least square discriminatory analysis (OPLSDA) model, as illustrated in Figure 1a and 1b.

**Figure 1.**
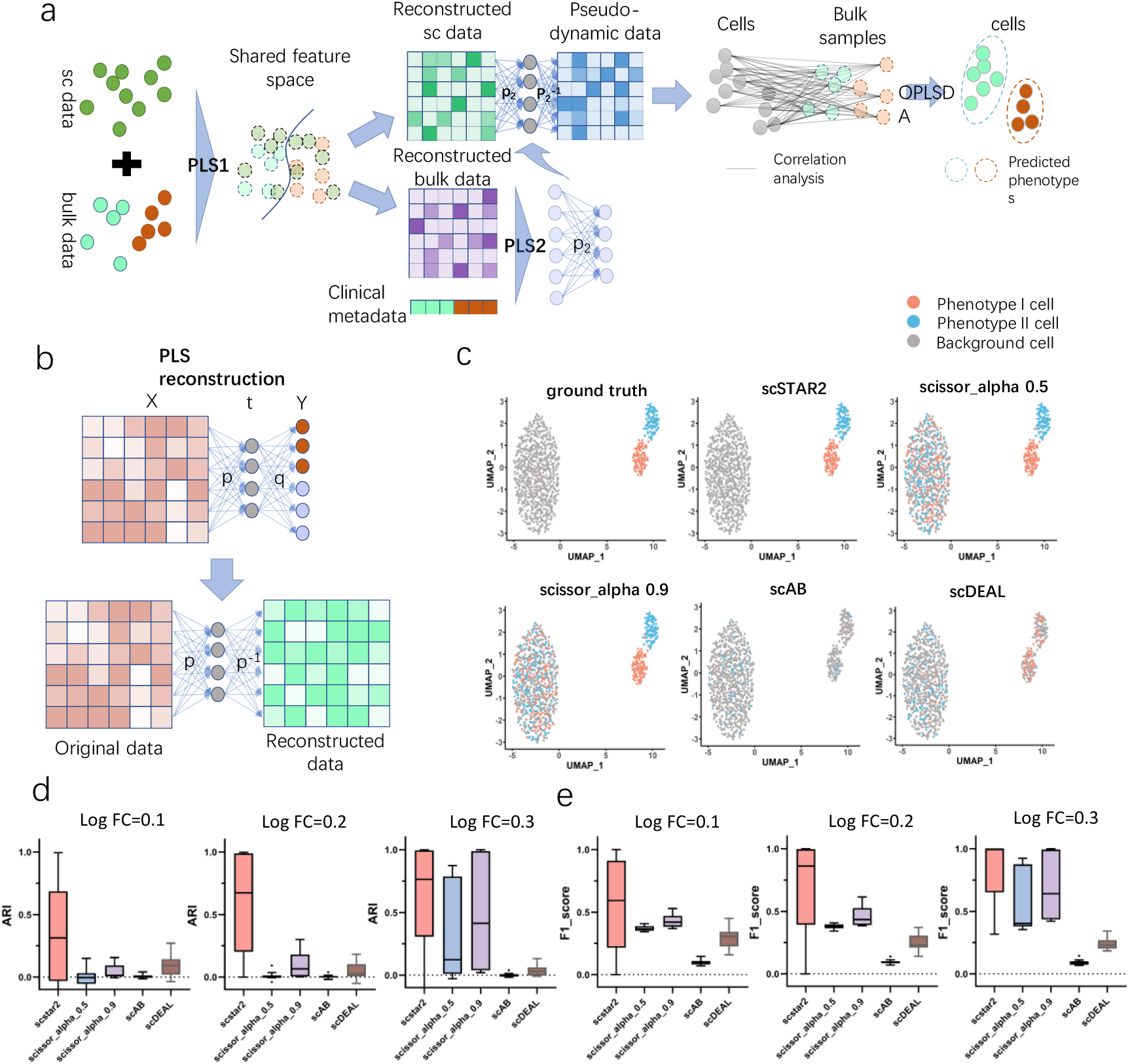
Schematic Workflow and Validation of scSTAR2 on Simulated Data. a)The diagram illustrates the scSTAR2 algorithm, detailing the process from data input through phenotype correlation. Initial inputs include a single-cell data matrix and a bulk RNA-seq expression matrix paired with phenotype annotations. b) These are processed through a partial least square (PLS) model to extract shared latent features and reconstruct data to emphasize phenotype-associated dynamics. c) The UMAP visualization demonstrates scSTAR2’s ability to align closely with ground truth in differentiating cell subpopulations by phenotype, outperforming other methods (scissor, scAB, scDlEAL, log FC=0.1), d) across varying levels of differential expression (DE) gene expression log fold changes (Log FC=0.1, 0.2, 0.3), as evidenced by higher ARI scores and F1 metrics.

### Evaluation of scSTAR2 on simulated data

To benchmark scSTAR2, we conducted evaluations against established methods such as scissor (with parameters alpha 0.5 and 0.9), scAB, and scDEAL, utilizing a range of simulated datasets. Our simulated datasets comprised bulk data, labeled with two distinct phenotypes, alongside corresponding single-cell datasets, ensuring a comprehensive evaluation setting. The singe-cell datasets contained one large subpopulation shared by both phenotypes and two small subpopulations unique to each phenotype, and the “unique to phenotype” cells were phenotype related cells (see more details in the Method section). In simulation, the average log fold change of differentially expressed (DE) gene expression is set as 0.1,0.2 and 0.3 to mimic low signal to noise ratio.

With the guidance of labeled simulation bulk datasets, each method was applied to identify the relationship between cells and phenotypes. The UMAP plots demonstrated scSTAR2’s consistency with ground truth (Figure 1c). The ARI results confirmed such observation which show that scSTAR2 achieved average values of 0.37, 0.59, and 0.77 in three simulation conditions, while the maximum of the other methods was less than 0.05, 0.1 and 0.5 (Figure 1d). Such results indicate that scSTAR2 can clarify more precise phenotype-related information in single cell populations. While scissor accurately identified true phenotype-related subpopulations in certain scenarios, it also incorrectly associated numerous background cells as related, highlighting a key area where scSTAR2 demonstrates its precision. Meanwhile, along with a larger log fold change of DE gene expression, most methods achieved better performances. Furthermore, the F1 score metric was used to illustrate the power of different methods in distinguishing the phenotype-related cells out of background cells. scSTAR2 achieved average F1 score of 0.54, 0.59 and 0.82, which is much higher than that of scissor (alpha 0.9, performed 0.43, 0.46 and 0.7) and other methods. In summary, scSTAR2 excels in discerning phenotype-related information within single-cell populations, effectively integrating this with the rich phenotype data from bulk samples, demonstrating its robustness and potential in single-cell data interpretation.

### scSTAR2 pinpoints heatshock Treg subtype by linking cellular profiles with patient prognosis

Expanding upon our previous identification of a unique Treg subtype with heatshock gene expression linked to patient prognosis [22], we sought to validate scSTAR2’s efficacy using scRNA-seq data from tumor tissues and survival-informed bulk RNA-seq data. This validation centered around recreating the findings of [22] by harnessing only tumor tissue-derived scRNA-seq data, supplemented by bulk RNA-seq data informed by survival metrics.

We integrated 1459 Treg cells from scRNA-seq HCC tumor samples [25], with 198 bulk RNA-seq specimens from the TCGA database (https://www.cancer.gov/tcga), stratified by survival rates to delineate good and poor prognoses (94 individuals with survival > 3 years categorized as good prognosis and 104 individuals with survival <1 year as poor prognosis). While initial categorization divided Treg cells into 5 clusters, scSTAR2’s processing further refined this to 6 distinct clusters (Figure 2a-b). A comparative analysis spotlighted variations in cell heterogeneities between the original data and those processed by scSTAR2 (Figure 2c). Notably, scSTAR2 earmarked the cluster scS_3 for its link to poor prognosis (Figure 2d)—a correlation affirmed by survival metrics generated via an online platform [26], leveraging the top 50 marker genes of scS_3 (Figure 2e). Elevated expression levels of heatshock Treg markers, HSPA5 and HSP90B1, were pinpointed in the Ori_3 and Ori_4 clusters, both of which converged into the poor prognosis-aligned scS_3 (Figures 2f-g, 2c). This underscored a potential defining feature of the poor prognosis Treg subtype: the pronounced expression of HSPA5 and HSP90B1, aligning perfectly with our previous findings in [22]. A mirrored pattern emerged in the LUAD dataset (Figures 2h-n).

**Figure 2.**
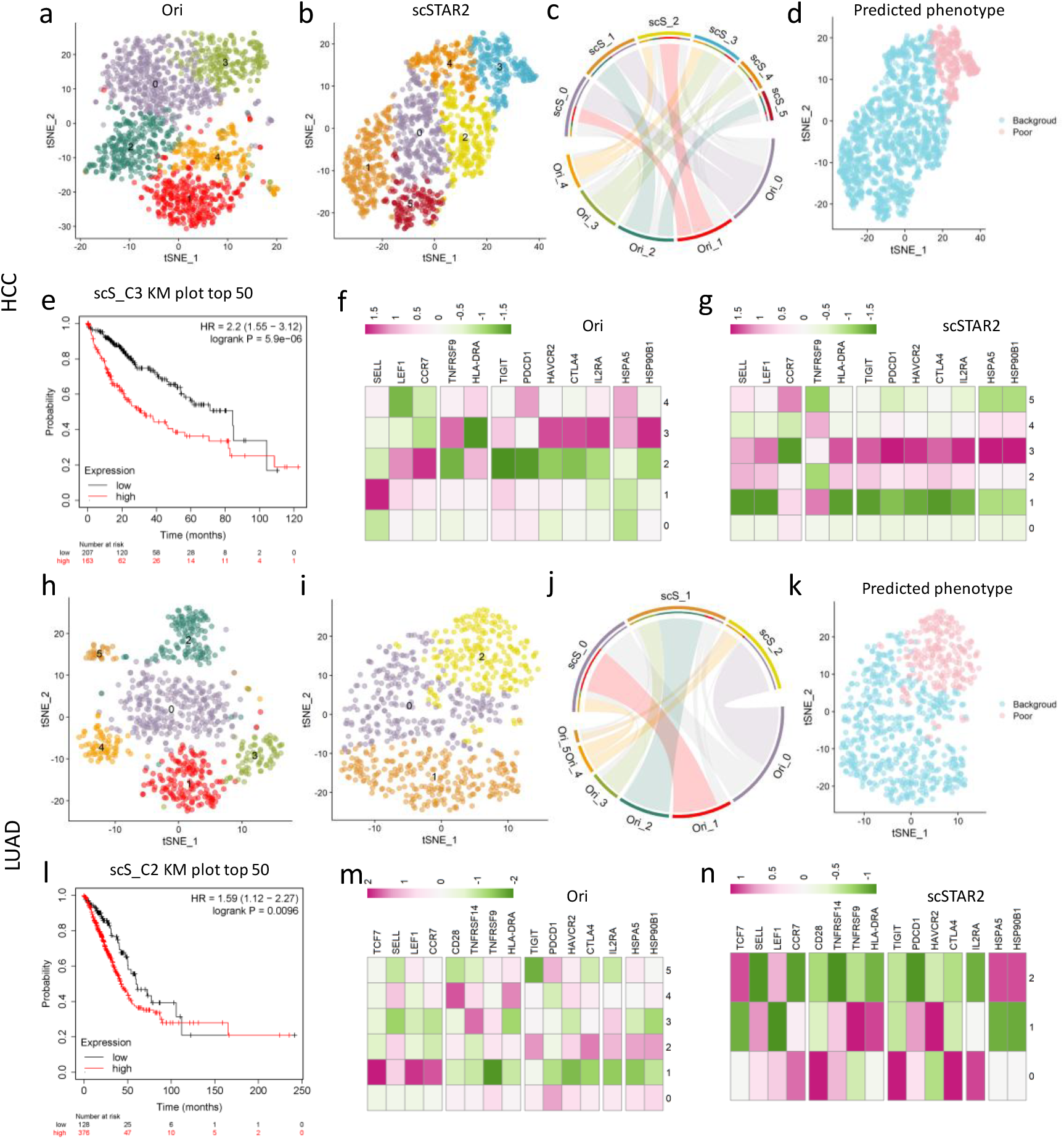
Heatshock Treg subtype discovery by scSTAR2. (a, b) baseline and post-process t-SNE plots showcasing cellular subdivisions for HCC. (c) Chord diagram elucidating the overlaps between original and scSTAR2-processed clusters. (d) Cellular phenotypic predictions derived from scSTAR2. (e) Kaplan–Meier overall survival curves of TCGA HCC patients with the top 50 genes of scSTAR2 cluster C3 (sorted by fold change), emphasizing the prognostic potential of scSTAR2’s categorizations. (f, g) Heatmaps of gene expression profiles highlighting crucial heatshock genes impacting patient outcomes in HCC. (h-n) The results obtained from LUAD data.

In conclusion, scSTAR2 proficiently uncovers clinically relevant cellular characteristics, enhancing the interpretive power of scRNA-seq data through the integration of phenotype information from bulk datasets.

### scSTAR2 unveils immune subtypes tied to heart failure by harmonizing single-cell and bulk transcriptomes

To further demonstrate the capability of scSTAR2 in seamlessly weaving scRNA-seq data with bulk RNA-seq and associated metadata, we exemplified scSTAR2 on scRAN-seq [28] and bulk RNA-seq data [29] from myocardial infarction heart failure (HF) studies. Collectively, 1121 B cells, 5069 CD8^+^ T cells, 9707 CD4^+^ T cells and 2505 NK cells were calibrated against 64 bulk samples (34 HF and 30 non-HF) using scSTAR2, respectively.

For CD8^+^ T cells, initial clustering identified 12 distinct clusters (Fig 3a). Post-scSTAR2 processing reduced these clusters to 5, suggesting extraneous, phenotype-indiscriminate cellular heterogeneities had been filtered out (Fig 3b). Linkages between the original and scSTAR2-modified clusters are mapped in Fig 2c. Notably, scSTAR2 earmarked cluster scS_3 for its strong association with HF (Fig 3d)—a claim corroborated when the marker genes of scS_3 manifested heightened expression in HF over non-HF samples (Fig 3e). Interestingly, it can be seen that new cell-cell variations which is invisible in the original data were revealed, e.g., the original cluster O_2 bifurcated, with a subset assimilating into the HF-linked cluster scS_3. Analogous patterns emerged across the other cell categories (Figure S1).

**Figure 3.**
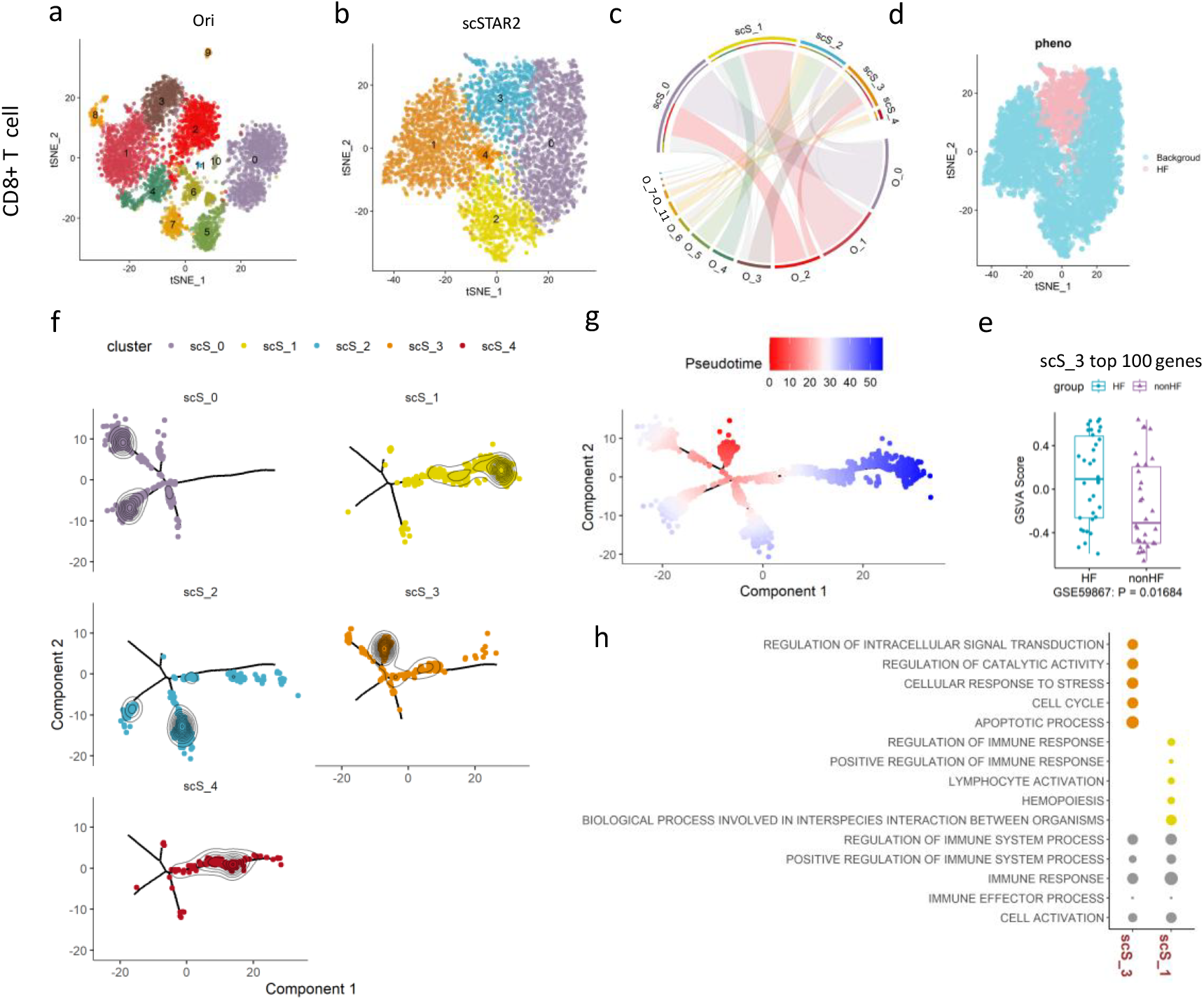
Heart failure associated CD8^+^ T cell characterization by scSTAR2. (a, b) baseline and post-process t-SNE plots showcasing cellular subdivisions of myocardial infarction heart failure patients. (c) Chord diagram representing transitions between the original and scSTAR2-processed cellular clusters. (d) Cellular phenotypic predictions derived from scSTAR2. (e) By Gene Set Variation Analysis (GSVA) analysis, the boxplot shows that the GSVA scores of top 100 maker genes of scS_3 are significantly higher in HF than nonHF patients. A two-sided Wilcoxon test was used to determine significance. (f) Pseudotime trajectory reconstructed on the scSTAR2 processed data, with estimated pseudotime scores of cells. (h) GO functions enriched in the cell clusters of interest.

Further characterization through pseudotime trajectory using monocle2 [1] analysis illuminated the HF-associated cellular dynamics within CD8^+^ T cells processed by scSTAR2 (Fig 3f). When use the scS_3 as the root state, scS_1 was referred to as a representation of non-HF characteristics (Fig 3g). Gene ontology (GO) assays revealed that HF-tied CD8^+^ T cells intensified apoptotic activities—aligning with prior studies [30], while the non-HF counterparts amplified immune responsiveness (Fig 3h).

We also applied scSTAR2 to analyze macrophage single-cell and bulk RNA-seq data in ovarian cancer patients undergoing chemotherapy (Figure S2). It identifies scS_3 (M1-like macrophages) as linked to treatment responders and scS_6 (M2-like macrophages) associated with non-responders (Figure S2 a, b and c). Marker genes from these clusters accurately predict chemotherapy outcomes in independent RNA-seq cohorts (Figure S2 d to g). Taken together, scSTAR2 forges a crucial link between phenotypic cues in bulk RNA-seq and single-cell details, unveiling cell characteristics pivotal to understanding disease pathogenesis.

### scSTAR2 maps prognostic insights onto spatial transcriptomic vistas

Spatial characteristics are pivotal in understanding cellular intricacies of multifaceted tissues. Recently developed spatially resolved transcriptomic technologies can profile the gene expression patterns in relation to spatial positioning, therefore, have great potential in disease studies [31-33]. However, current spatial transcriptomic (ST) data interpretation strategies often lean on pre-established knowledge. For example, the discernment of an immunosuppressive microenvironment at tumor borders is based on the congregation of M2 macrophages and exhausted T cells [34]. Such methodologies, although informative for recognized characteristics, are somewhat blinkered when it comes to unveiling novel clinical phenotype-linked spatial attributes. To the best knowledge of the authors, tools explicitly linking spatial attributes to clinical phenotypes remain in their infancy.

We demonstrate scSTAR2’s utility in deriving insights from ST data of a breast cancer patient [31], correlating spatial gene expression with clinical phenotypes. The ST data encompassing 613 spots were reconstructed with the guidance of breast cancer bulk RNA-seq data from TCGA database. The bulk RNA-seq dataset contains 923 samples with 438 good prognoses (survival time >3 years) and 485 poor prognoses (survival time <2 years). The reconstructed ST data were clustered into 6 clusters (Figure 5a), among which the clusters 0, 3, 4, 5 were assigned as good prognosis associated and the cluster 2 was poor prognosis by scSTAR2 (Figure 5b).

**Figure 5.**
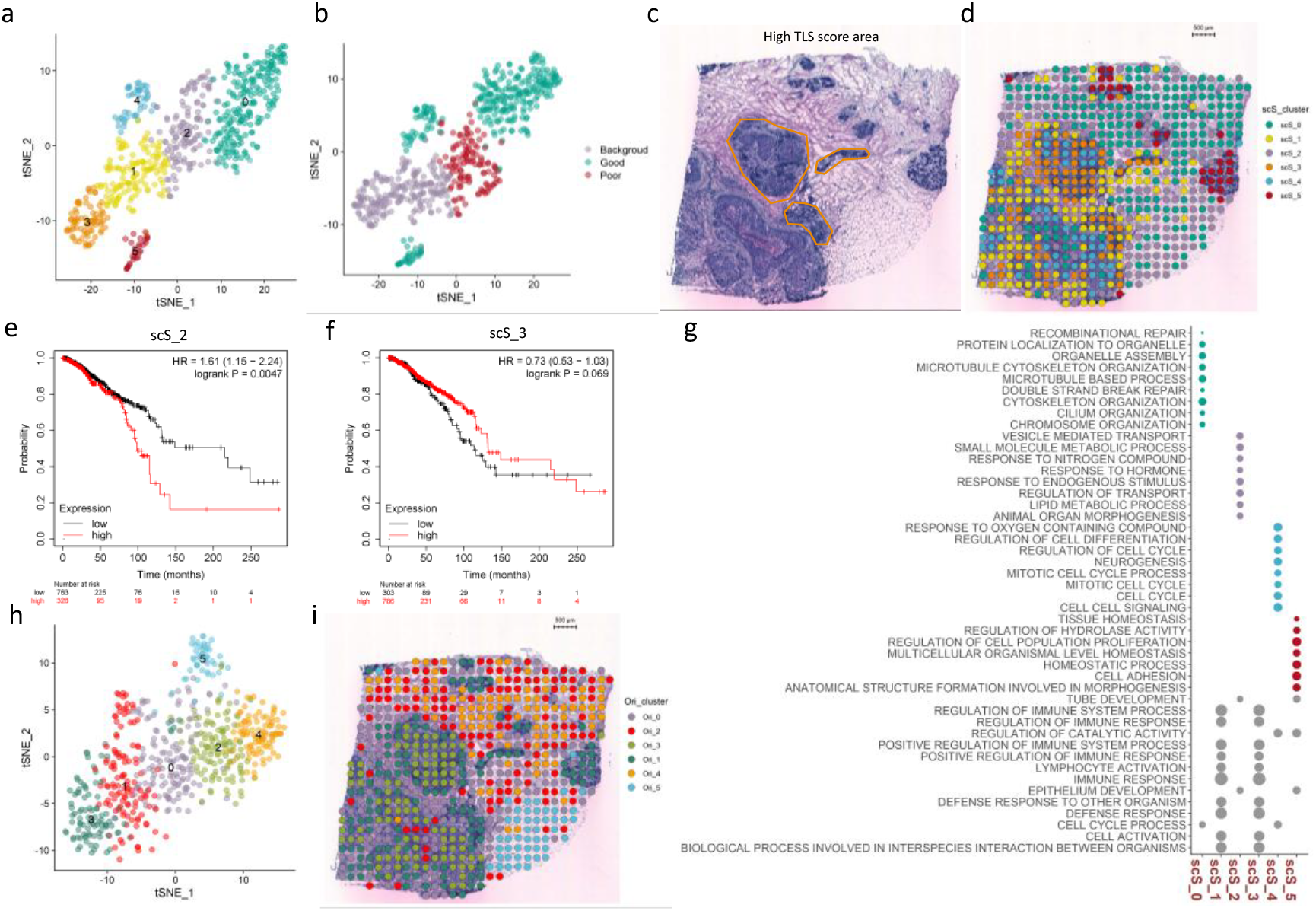
Breast cancer clinical phenotypes associated spatial regions identification on spatial transcriptomic data by scSTAR2. (a, b) t-SNE plots demonstrating cellular diversity and their subsequent phenotypic groupings. (c) Histological representation indicating areas with TLS region highlighted. (d) The spatial mapping of spots with color coded cluster indexes. (e, f) Kaplan-Meier survival curves showing the association between gene expression and patient survival for two clusters scS_2 and scS_3. (g) Dot plot detailing the association of key gene oncology biological processes with specific cell groups, where the size indicates the significance of genes involved. (h, i) Spots clustering results and their spatial mappings on original gene expression spectrum.

Original Tertiary lymphoid structures (TLSs) identifications [31] (Figure 5c) were refined by scSTAR, suggesting a broader spatial distribution of TLS-associated cells beyond previously recognized regions. Cluster 3 covered the TLS regions annotated in the original study. Cluster 1 surrounded around the annotated TLS regions (Figure 5d) and had the identical molecular functions as Cluster 3 (Figure 5g). This observation implies that both Cluster 1 and 3 should be TLS. Such results indicate that scSTAR2 has higher sensitivity in dissecting ST data than conventional methods. However, scSTAR2 did not associate the TLS regions with significant prognosis, which was confirmed by using an independent webserver [26] (Figure 5f).

Noticeable, the poor prognosis associated Cluster 2 mainly corresponded to adipocytes which indicates it might be cancer-associated adipocytes [35]. The observation was validated using an independent webserver [26] (Figure 5e). GO analyses of the marker genes of different clusters indicate that poor prognosis associated Cluster 2 had active lipid metabolism function, which is consistent to previous studies [36]. However, directly analyzing the ST data failed to separate TLSs regions (Figure 5h and 5i).

Comparatively, scissor, scAB, and scDEAL displayed less specificity in spatial spot identification (Figure S2). It can be seen that scissor identified spots almost randomly scattered across the spatial chip, which were less informative (Figure S2 b and d). scAB only identified the TLS and some para-tumor areas as phenotype associated but most of the other regions were annotated as background (Figure S2f). scDEAL identified a TLS area as poor prognosis related (Figure S2h). Those results were all contradictory to some extent with existing knowledge.

Further validation of scSTAR2, using ST data from an ovarian cancer patient (https://www.10xgenomics.com/resources/datasets/human-ovarian-cancer-1-standard) together with the TCGA ovarian RNA-seq data, among which 77 samples with survival time >5 years and 75 samples with survival time <1 year, confirmed its efficacy in associating spatial spots with survival outcomes. Our analyses identified a cluster of spatial spots associated with long survival phenotype as illustrated in Figure S3. Such observation was confirmed by the webserver [26] (Figure S3d).

In conclusion, scSTAR2 enhances spatial transcriptomic interpretation by integrating bulk RNA-seq with clinical metadata, offering a powerful tool for disease pathogenesis studies

### scSTAR2 identifies immune abundant area associated with TLS in ST data

TLSs are found in many cancers and their prognostic and predictive value in mediating antitumor immunity has spurred increased interest in their role [37, 38]. Here, we demonstrated how to use scSTAR to highlight the TLSs from ST data by incorporating independent ST data with annotated TLS areas.

We collected 10x visium ST data of renal cell carcinoma with manual annotation of TLS [38]. We acquired four FFPE slides from the dataset and utilized scSTAR2 in combination with cellrank2 [39] to analyze the ST data from renal cell carcinoma, leveraging pre-annotated TLS and no_TLS areas as references. Utilizing one single ST slide with TLS area manually annotated as reference, we seek to find TLS associated area in other three ST slides. Consequently, scSTAR2 predicted TLS-associated areas in the three query slides, showing high overlap with the manual annotations and extending to adjacent regions suggestive of nascent TLS formation (Figure 6 a-c). By combining the TLS spots from the reference slide and all ST spots from the three query slides, we observed that the scSTAR2 predicted TLS spots gathered together with the reference TLS spots (Figure 6 d-f). Cellrank2 analysis revealed a developmental trajectory towards annotated TLSs within these predicted areas, based on diffusion pseudotime [40], which is a feature absent in the raw ST data (Figure6 g-i, Figure S4 a-i). This prediction was supported by the high probability of these areas developing into TLSs, as determined by analysis of terminal states and fate probabilities (Figure6 j-l), and the strong correlation between predicted TLS areas and the expression of known TLS marker genes, such as CD3D, CD4, CD8A, CD79A (Figure S5-S7, Supplementary Table 1).

**Figure 6.**
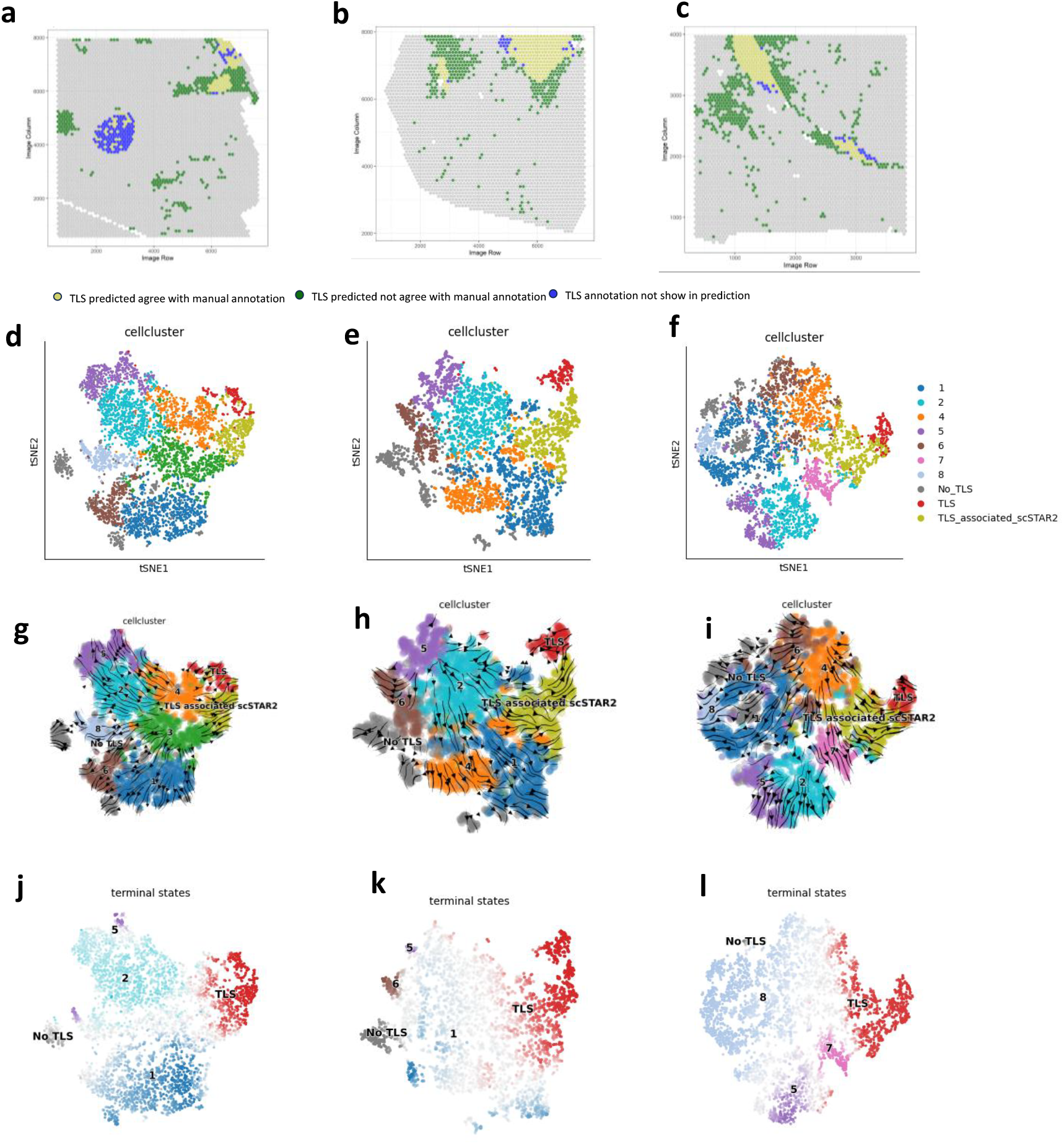
scSTAR2 highlights TLS features by incorporating pre-annotated ST data. (a-c), the results of the predicted TLS area by scSTAR2 are overlapped with manual-annotation, slide 1-3. The yellow color represents the spot both annotated as TLS by scSTAR2 and manual annotation; green represents the spot annotated as TLS by scSTAR2 but not manual annotation; bule represents the spot annotated as TLS by manual annotation but not scstar2. (d-f), the tSNE cluster results of query data and reference data after scSTAR2. TLS and No_TLS clusters are spots from reference data, TLS_associated_scSTAR2 represent spots assigned as TLS associated by scSTAR2. (g-i), the streamline results by cellrank2, based on diffusion pseudo-time data after processed with scSTAR2. (j-l), the fate probabilities of each spot, each color means the spot is most likely to develop into the corresponding terminal states assigned by cellrank2.

These findings demonstrate scSTAR2’s ability to identify TLS-associated regions with developmental potential, exceeding the capabilities of standard ST analysis.

### scSTAR2 associates patients’ prognosis with scATAC-seq data

scSTAR2’s adeptness in multiomics integration is further exemplified by processing scATAC-seq data collected from 5 ccRCC patients, RCC81, RCC84, RCC86, RCC87, RCC94 in [41] aligning chromatin accessibility profiles with clinical prognoses from TCGA-ccRCC. We used 152 bulk RNA-seq samples with OS (overall survival)>5 years as good prognosis, and 93 samples with OS<1 year as poor prognosis. First, 10740 cells were collected from the matched samples and profiled using scRNA-seq. The cell types were annotated based on the canonical marker reported in the original study (Figure S8a). Second, the scATA-seq peaks were transfer to gene activity matrix using Seurat function ‘GeneActivity’. Third, the cell types of scATAC-seq data were inferred by Seurat function ‘TransferData’ on scRNA-seq data (Figure S8b).

Focusing on 5717 annotated CD8^+^ T cells from the scATAC-seq dataset, conventional analysis revealed 8 clusters, (Figure 7a), whereas scSTAR2 processing identified a subgroup of CD8+ T cells, scS_1 (Figure 7b), with good prognosis (Figure 7c). Survival analysis using the genes identified by the Seurat “FindAllMarkers” function (top 30 largest fold changes) confirmed the finding of scSTAR2 (Figure 7d). The cluster scS_1 was predominantly from the cluster Ori_3 (Figure 7e). The top 10 markers of scS_1 encompassed AGXT2 and CLDN2, both of which showcased pronounced chromatin accessibility in Ori_3 (Figure 7f and 7g) and were reported to be associated with favorable kidney cancer patient survival [42, 43]. However, the previous studies used conventional methods and lost the information of single cells, therefore, failed to allocate the expression patterns of the two key genes on tumor microenvironment cells. Our findings not only extend current knowledge but also underscore the critical role of microenvironment cells in the tumor landscape, offering new insights into kidney cancer patient survival.

**Figure 7.**
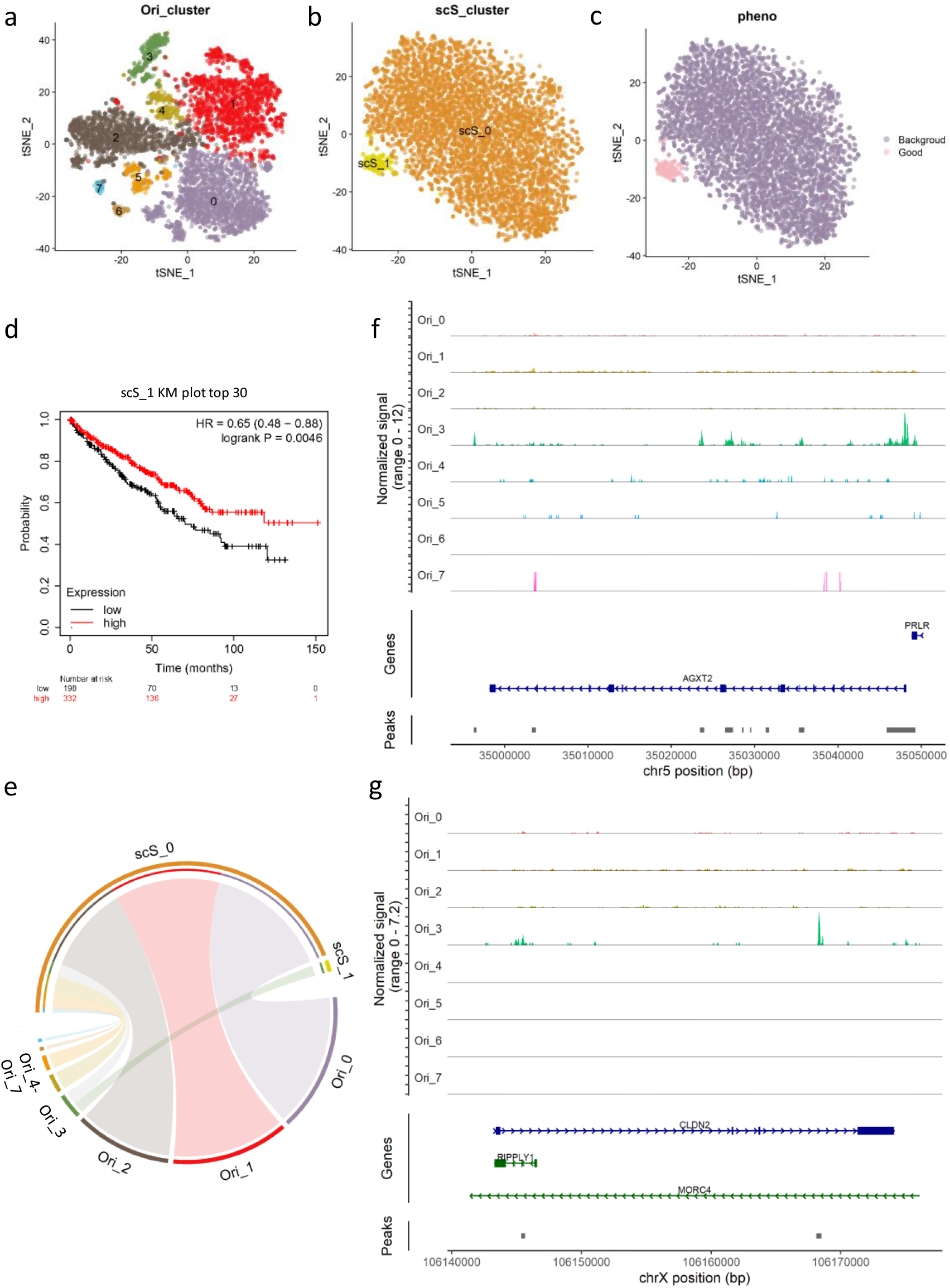
Association of scATAC-seq data with tumor prognosis phenotypes using scSTAR2. (a-c) Cell subtype phenotypic predictions derived from original and scSTAR2 processed gene activity data. (d) Kaplan-Meier survival curves showing the association between gene expression and patient survival for scS_1. (e) Chord diagram representing transitions between the scSTAR2-processed cellular clusters and original clusters. (f, g) AGXT2 and CLDN2 chromatin accessibility profiles of each cluster.

In summary, scSTAR2 proves itself as a powerful tool in revealing clinically relevant chromatin accessibility features, affirming its utility in multiomics data integration.

## Discussion

scSTAR2 adeptly bridges wealthy clinical metadata to single cells to decipher critical disease associations. Its capability to extract detailed patterns across various datasets, as demonstrated through heart failure scRNA-seq data, cancer scRNA-seq data, spatial transcriptomics, and scATAC-seq data, reinforces its potential as a versatile multiomics integration tool.

As indicated by ARI and F1 score, scSTAR2 is more robust than existing methods. This improvement is largely attributed to scSTAR2’s unique reconstruction strategy, which filters out gene expression variations irrelevant to the phenotype, enhancing the identification of phenotype-specific cells. Existing methods, like scissor, scDEAL and scAB, identified phenotype-specific cells directly from the original fuzzy single-cell data. However, scSTAR2 first reconstructs the single-cell data with the guidance of phenotype information. Such procedure removes all gene expression variations irrelevant to phenotypes, which improves the reliability of phenotype-specific cell identification.

When applied to ST data, scSTAR2’s reconstruction procedure unveiled expanded TLS structures, suggesting a broader spatial involvement than previously recognized, enhancing our understanding of tissue microenvironments. When applied on ST data, a group of spots, i.e., cluster 1 in Figure 5a, was found highly to have almost identical molecular functions as TLS regions and these spots also spatially surrounded the annotated TLS structures. Such results implied that TLS structures may be more expanded than they were usually identified from IHC images. Such deduction was also supported by the results illustrated in Figure 6. scSTAR2 may facilitate ST data interpretation by automatically and readily pinpointing the clinical phenotype related tissue regions.

scSTAR2 extended the application scenarios of single cell techniques and improved the interpretability of single cell results by reconstructing single cell data in a novel way. We believe the development of scSTAR2 may promote the clinical applications of single cell techniques. Furthermore, we also realized that the performance of scSTAR2 may be compromised in dealing with larger and more complex single cell datasets due to machine learning models. In the future works, deep learning models may be applied in place of machine learning models, which can cope with large datasets and complicated linear and nonlinear relationship between multiomic data.

## Methods

### Gene filtering

To map clinical phenotype information on individual cells or spatial spots, scSTAR2 first integrates single cell data with bulk RNA-seq data (referred to as RNA-seq). Before integrating the two types of data, gene filtering is performed for two purposes. 1) reduce the random noise interferences, especially in scRNA-seq data. 2) RNA-seq data contain all types of cells, however, we process one cell type from scRNA-seq data each time. The components of the untargeted cell types in RNA-seq may cause unwanted interferences due to limited specificity of marker gene expression. To address this problem, we applied OGFSC [44] to scRNA-seq data and eliminated genes with variances below the noise threshold curve determined by the α parameter. To maintain maximum signal integrity, the value of α is fixed to be 0.5. Then the shared genes between the OGFSC selected ones and those annotated in RNA-seq data are used.

### Latent feature extraction

Given the RNA-seq data, the corresponding clinical phenotype information of interest as reference, and the scRNA-seq data, the shared biological characteristics across the scRNA-seq and RNA-seq data are extracted. Let’s denote the scRNA-seq data as **X**_***S*** = {_***x***_*S*1_, ***x***_*S*2_, …, ***x***_*Sn*1_} and the RNA-seq data as **X**_***B*** = {_***x***_*B*1_, ***x***_*B*2_, …, ***x***_*Bn*2_}, where ***x***_*Si*_, ***x***_*Bj*_ ∈ ℝ^*N*^ representing the cell ***S***_***i***_ from scRNA-seq data and the sample ***B***_***j***_ from RNA-seq data. *N* denotes the number of shared genes, and *n*_1_ and *n*_2_ denotes the number of cells and samples, respectively.

Since the two types of data aim to characterize the same problem, it is reasonable to assume that there exist low dimensional feature spaces, in which the projections of the scRNA-seq and RNA-seq data are similar. At the same time, the configuration of the original data can be preserved to the maximum extent in the feature space. To fulfill the above requirements, we construct a contrast function:

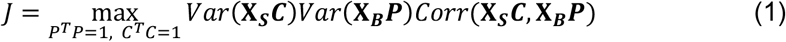

Assume we want to reduce the dimensionality of X_*S*_ by orthonormal transformation *C*, and we want to keep the squared reconstruction error small,

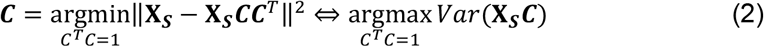

‖. ‖ denotes L2 norm. Therefore, the first two terms in Eq. 1 guarantee the constructed transformation matrix ***C*** = {***c***_1_, ***c***_2_, …, ***c***_*n*_} and **P** = {***p***_1_, ***p***_2_, …, ***p***_*n*_} best preserve the original configurations of **X**_***S***_ and **X**_***B***_. The last term *Corr*(**X**_***S***_***C*, X**_***B***_***P***) guarantees the projected **X**_***S***_ and **X**_***B***_ have maximum correlation. By making a slight modification of Eq. 1, we have,

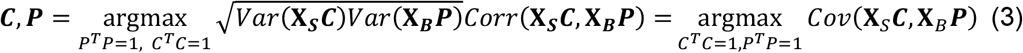

which is the contrast function of partial least square (PLS) model. A widely applied solution of Eq. 3 is nonlinear iterative partial least squares (NIPALS). Using NIPALS, the first component of transformation matrix **c**_1_ and **p**_1_ can be calculated as,

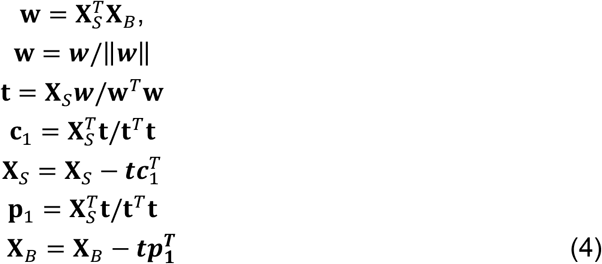

Repeat the above procedure for n times to estimate ***C*** and **P**. Let 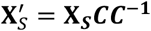 and 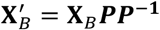 the reconstructed scRNA-seq and RNA-seq data containing only the desired features, which are the best representations of **X**_*S*_ and **X**_*B*_, and are mutually highly correlated.

### Pseudo-dynamical space reconstruction between two phenotype conditions

Once the latent features 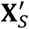 and 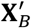 are extracted, a PLSDA (partial least square discriminatory analysis) is applied to project the scRNA-seq data into another feature space. The new feature space should be spanned on the dynamical features associated with the discrimination of different phenotypes, and therefore be referred to as pseudo-dynamical space. The pseudo-dynamical space is designed to represent single-cell-resolution gene expression dynamics between different phenotypes specified by the clinical metadata associated with RNA-seq datasets. In scSTAR2, the pseudo-dynamical space projection model is constructed based on 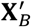 and clinical metadata, then the obtained model is applied on 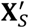.

It is reasonable to assume the meta information of each sample (patient or animal) are related to its RNA-seq data 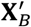. Take two-phenotype-conditions as an example, a dummy matrix Y with dimensions *n*_2_ × 2 is constructed, which contains binary values. The two columns indicate the phenotype condition each sample belongs to, as denoted by (1, 0) or (0, 1). By optimizing the covariance based contrast function using NIPALS method,

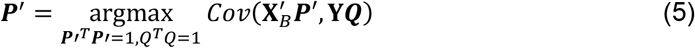

we obtain a new transformation matrix ***P***^′^. The space spanned by ***P***^′^ represents the features which only associated with the discrimination of two phenotype conditions. The RNA-seq data can be projected in the pseudo-dynamic space by 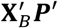.

Considering the common characteristics shared between 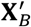 and 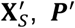, ***P***^′^ can also be used to reconstruct 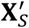 from the pseudo-dynamic space

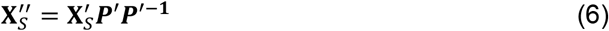

### Association of single cells with phenotype

Theoretically, 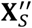 is dominated with the cell type-specific gene expression features associated with the dynamics between the two phenotype conditions. However, it is insufficient to determine the association between specific cells and phenotype conditions by 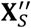 self. To tackle this issue, scSTAR2 first clusters the cells based on 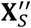 to characterize the phenotype associated heterogeneous cell subpopulations. Then, by evaluating the correlation between cell clusters and phenotype-annotated RNA-seq data, we could associate the cell clusters with phenotypes.

The cells 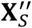 are clustered using Seurat R package. The genes specifically highly expressed in each cluster are obtained using ‘FindAllMarkers’ function of Seurat. As in reconstructed scRNA-seq data, all components irrelevant to the discrimination of different phenotypes have been eliminated, the high expression markers identified from each cell cluster represent the contribution of these cells to the discrimination of phenotype conditions. To affirm it, a Pearson correlation matrix 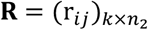 is calculated between each cell from a cluster and a bulk sample on the identified marker genes to quantify the similarity between cells and bulk samples, where *k* denotes the number of cells in the cluster.

A reasonable assumption is that if a cluster of cells has higher correlation with the bulk samples from one phenotype than the other, this cluster of cells may be associated with the phenotype. To test the significance of the similarity, we adopted OPLSDA (orthogonal partial least square discrimination analysis) [45], which is an extension version of PLSDA.

Here, we search for the common features of the cell-sample correlation matrix **R** and a dummy matrix **Y** indicating the phenotypes each sample belongs to. Compared to standard PLSDA, OPLSDA includes an orthogonal filtering step to condense all discriminatory features into the first PLS component, which enables us to extracted the desired information by interrogating the first PLS component only. The OPLSDA model between **R** and **Y** can be calculated as,

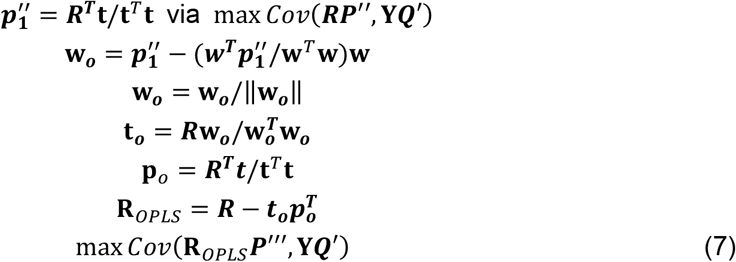

To identify the cell-phenotype associations, the correlation r_*i*−*OPLS*_ between 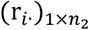 and the first OPLSDA score **t**_***OPLS***,**1**_ are evaluated. A significant r_*i*−*OPLS*_ (p-value<0.05) indicates that the cell *i* is more similar to one phenotype than the other. The cell *i* is associated with phenotype A if r_*ij*(*j*∈*phenotype A*)>_r_*ij*(*j*∈*phenotype B*)_ and 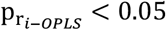.

Furthermore, a cross-validation Q^2^ statistic is calculated to evaluate the discriminatory of the correlation values between the two phenotypes. The OPLSDA model is considered valid if the model actual Q^2^ is significantly higher than the Q^2^ obtained by permutation test [46].

We define that if the cells from a cluster have significant and consistent association with a phenotype and at the same time the p-value of model Q^2^ is less than 0.05, the whole cluster of cells is associated with the phenotype.

### Simulate datasets

Similar with that in Scissor, we generated simulation datasets containing 1000 cells and 5000 expression genes with Splatter [47]. The cells were from three simulated cell types with proportions of 0.8, 0.1 and 0.1. The proportions of differentially expressed genes between each group and the others were 0.1, 0.01 and 0.01. Then, we assigned the three groups of cells into two phenotypes. Respectively, the large group of cells were set as background cells that were shared by both phenotypes, while the other two groups were set as unique to each phenotype. To simulate the bulk data of each phenotype, we randomly sampled 1000 cells (replacement set as Ture) within the cells belonged to each phenotype and take the average gene expression as bulk gene expression profile. Utilizing different random seeds, we generated 50 bulk samples of each phenotype.

To investigate the performance of scSTAR2 and other algorithms in dealing with datasets of different divergence, we simulated datasets with the parameter “de.facLoc” set as 0.1, 0.2 and 0.3 indicting different log fold changes (FC), where the larger log FC represents larger divergence between the cell groups. Taking the previous simulation as one simulation instance, we generated 10 instances under each set of parameters.

### Evaluation metrics

#### ARI

The group results (positive-related, negative-related and non-related) of cell populations obtained by different methods on the simulated data were evaluated by the adjusted Rand index (ARI)

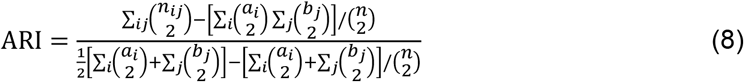

where *n*_*ij*_ are contingency table entry values and *a*_*i*_ and *b*_*j*_ are the sums of the *i*th row and the *j*th column of the contingency table, respectively. The closer the ARI value is to 1, the closer it is to the true cluster.

#### F1 score

The F1 score is a metric commonly used in binary classification to evaluate the model’s performance. Here, we use it to evaluate the performance of distinguishing phenotype-related cells from non-related (background) cells of methods.

Precision represents the accuracy of the positive predictions made by the methods.

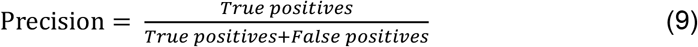

Recall is also called sensitivity or true positive rate. It represents the ability of the model to capture all the positive instances.

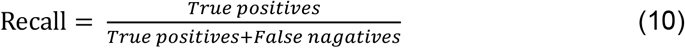

The F1 score is then calculated as the harmonic mean of precision and recall:

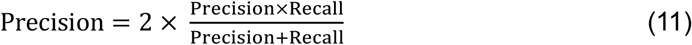

The F1 score ranges from 0 to 1, where a higher value indicates better performance.

### Experimental Datasets

To illustrate the capability of scSTAR2 in discovering heatshock Treg subtype, a human hepatocellular cell carcinoma tissue dataset (GSE140228) and a human lung squamous cell carcinoma tissue dataset (GSE99254) were adopted in together with the corresponding TCGA datasets. Validation of scSTAR2 in non-cancer investigations by revealing heart failure associated cellular features was adopted from a scRNA-seq dataset collected from post-myocardial infarction heart failure patients (GSE145154) and a bulk study containing post-myocardial infarction heart failure and non-heart failure patients (GSE59867). An ovarian cancer chemotherapy single cell dataset (GSE165897) and a bulk dataset (GSE156699) were applied to validate the potential of scSTAR2 in precision cancer treatment applications. The spatial transcriptomics data of renal cell carcinoma (RCC) with TLS annotation can be accessed by GSE175540. The spatial transcriptomics data from breast cancer (https://github.com/almaan/her2st) and ovarian cancer (https://www.10xgenomics.com/resources/datasets/human-ovarian-cancer-1-standard) were integrated with the corresponding TCGA bulk RNA-seq data to demonstrate the capability of scSTAR2 in combing clinical phenotypes with spatial features. The scSTAC-seq data (GSE207493) was adopted to demonstrate the versatility of scSTAR2 in dealing with multiomics data.

## Author Contributions

X.Z., J.H., and WT.C. conceived the study and the experimental setup; AZ. Z. made the scSTAR2 R package; X.Z., JW.Z., FL.D., AZ. Z., and YJ.L. performed the data analysis and wrote the manuscript; LJ.T., XB.S. and H.H.Y.T. assisted with the review and editing of the draft.

## Code availability

The scSTAR2 original code and full tutorials are available at https://github.com/Hao-Zou-lab/scSTAR2.

## KeyPoints

- scSTAR2 is an algorithm that integrates multi-omics data to identify cell signatures associated with clinical phenotypes, overcoming limitations of traditional methods.
- By reconstructing scRNA-seq and ST and leveraging phenotype information from independent data, scSTAR2 reduces interference from irrelevant noise, enabling more accurate identification of cell subpopulations and spatial regions related to clinical phenotypes.
- The algorithm has been successfully applied to various single-cell omics data (including scRNA-seq, scATAC-seq, and spatial transcriptomics), revealing cell subtypes and tissue structures associated with patient prognosis, such as heat-shock Treg subtypes and tertiary lymphoid structures.

