## Supplementary Figures for "scSTAR2: a multiomics integration algorithm to reveal disease-specific cellular signatures by bridging single-cell resolution features and clinical metadata"

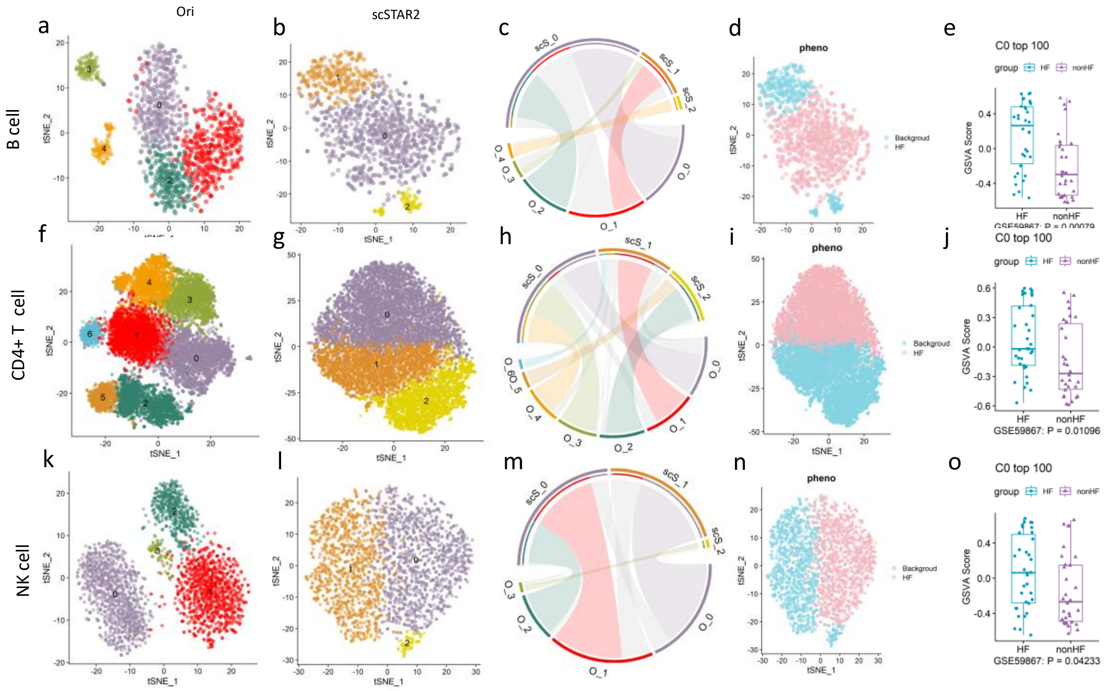


**Figure S1.** Heart failure associated immune cell characterization by scSTAR2. (a, b, f, g, k, l) baseline and post-process t-SNE plots showcasing cellular subdivisions of myocardial infarction heart failure patients. (c, h, m) Chord diagram representing transitions between the original and scSTAR2-processed cellular clusters. (d, i, n) Cellular phenotypic predictions derived from scSTAR2. (e, j, o) Validation of scSTAR2 identified cell-phenotype association on bulk data.

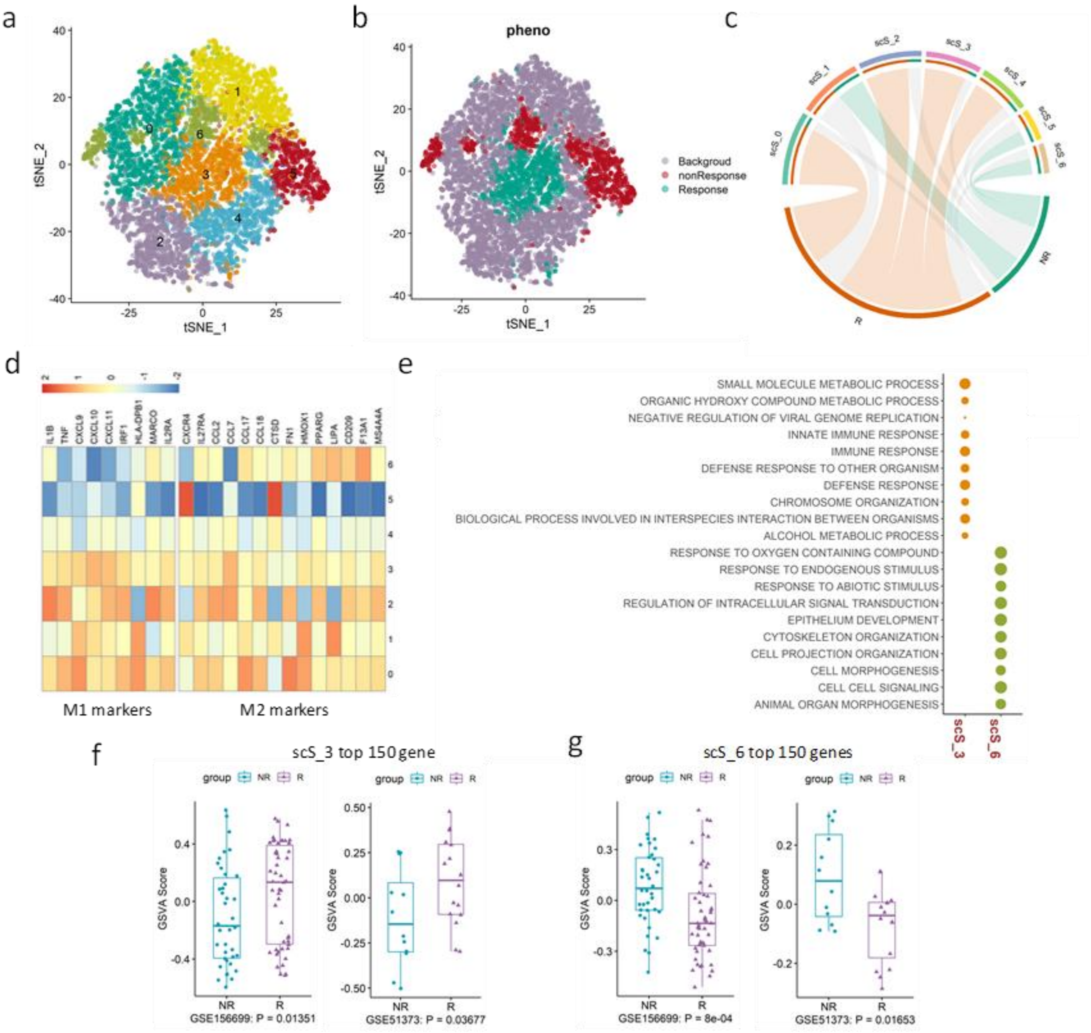


**Figure S2**. Chemotherapy outcome pattern associated macrophage subtypes identification by scSTAR2. (a, b) Cell subtype phenotypic predictions derived from scSTAR2. (c) Chord diagram representing transitions between the scSTAR2-processed cellular clusters and response pattern categories. (d) Heatmap of M1 and M2 marker gene expression across various cell subtypes, with color intensity denoting expression levels. (e) Dot plot detailing the association of key gene oncology biological processes with specific cell groups, where the size indicates the significance of genes involved. (f, g) Boxplots show that the GSVA scores of maker genes of scS_3 and scS_6 are significantly different between chemotherapy responders and non-responders. A two-sided Wilcoxon test was used to determine significance.

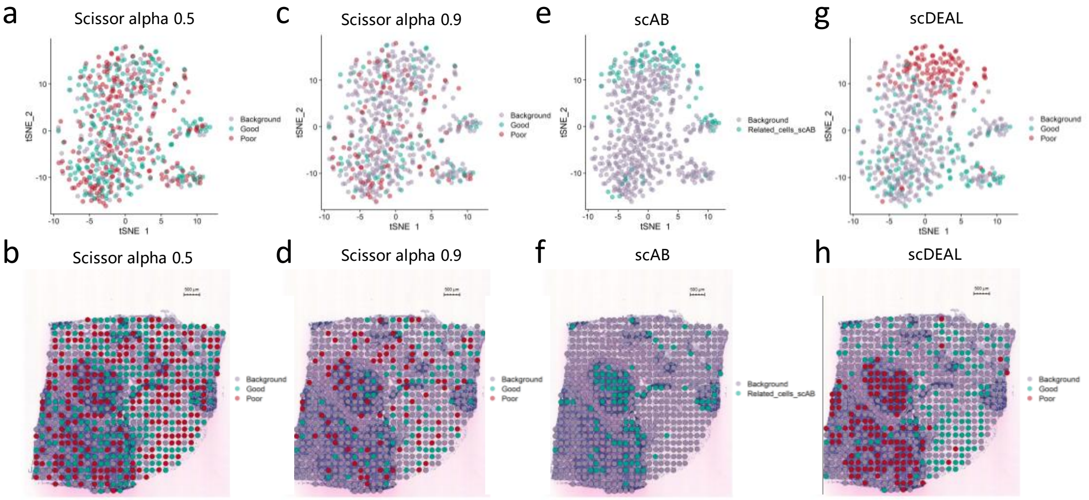


**Figure S3.** Breast cancer clinical phenotypes associated spatial regions identification on spatial transcriptomic data. Spots clustering results and their spatial mappings on original gene expression spectrum by scissor (a-d), scab (e, f) and scDEAL (g, h).

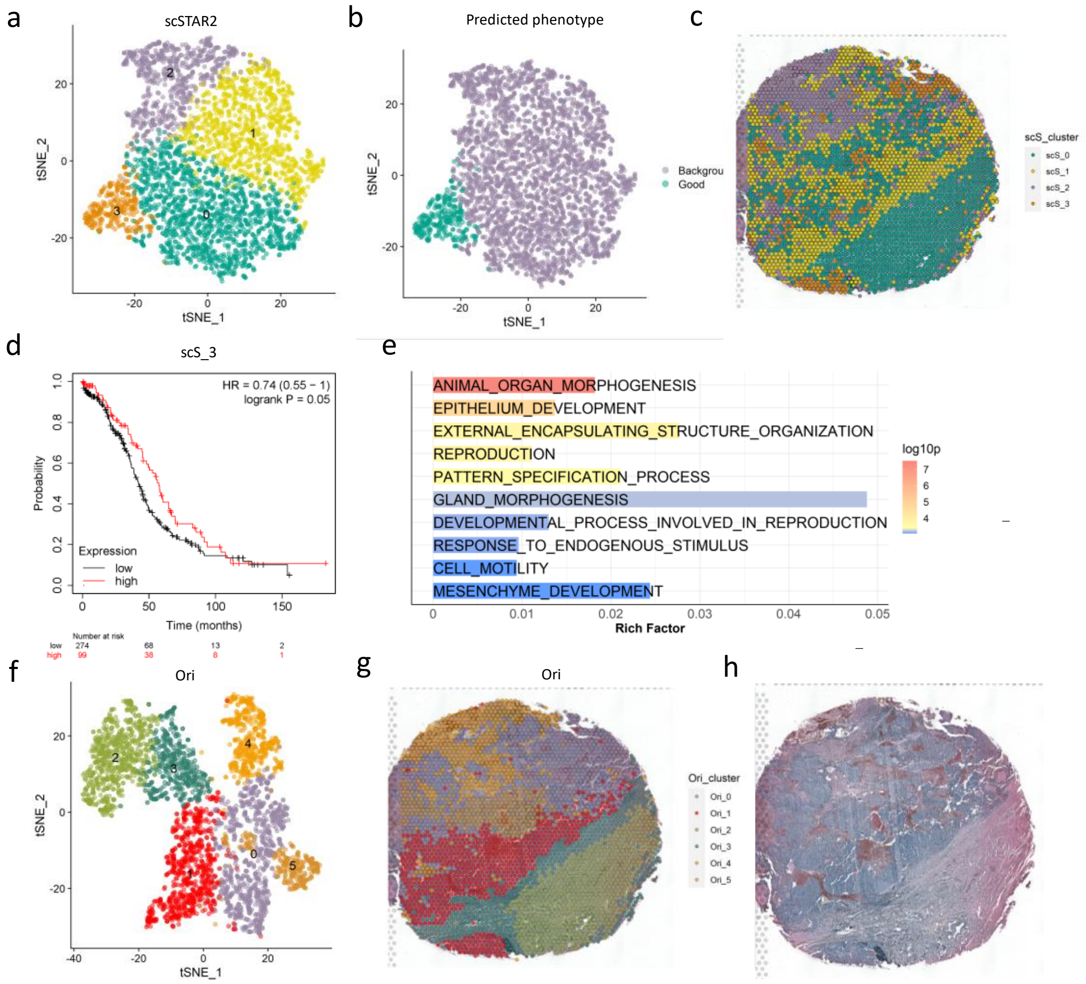


**Figure S4.** Ovarian cancer clinical phenotypes associated spatial regions identification on spatial transcriptomic data by scSTAR2. (a, b) t-SNE plots demonstrating cellular diversity and their subsequent phenotypic groupings. (c) The spatial mapping of spots with color coded cluster indexes. (d) Kaplan-Meier survival curves showing the association between gene expression and patient survival for scS_3. (e) The enriched gene oncology biological processes associated with scS_3. (f, g) Spots clustering results and their spatial mappings on original gene expression spectrum. (h) Histological representation of the sample.

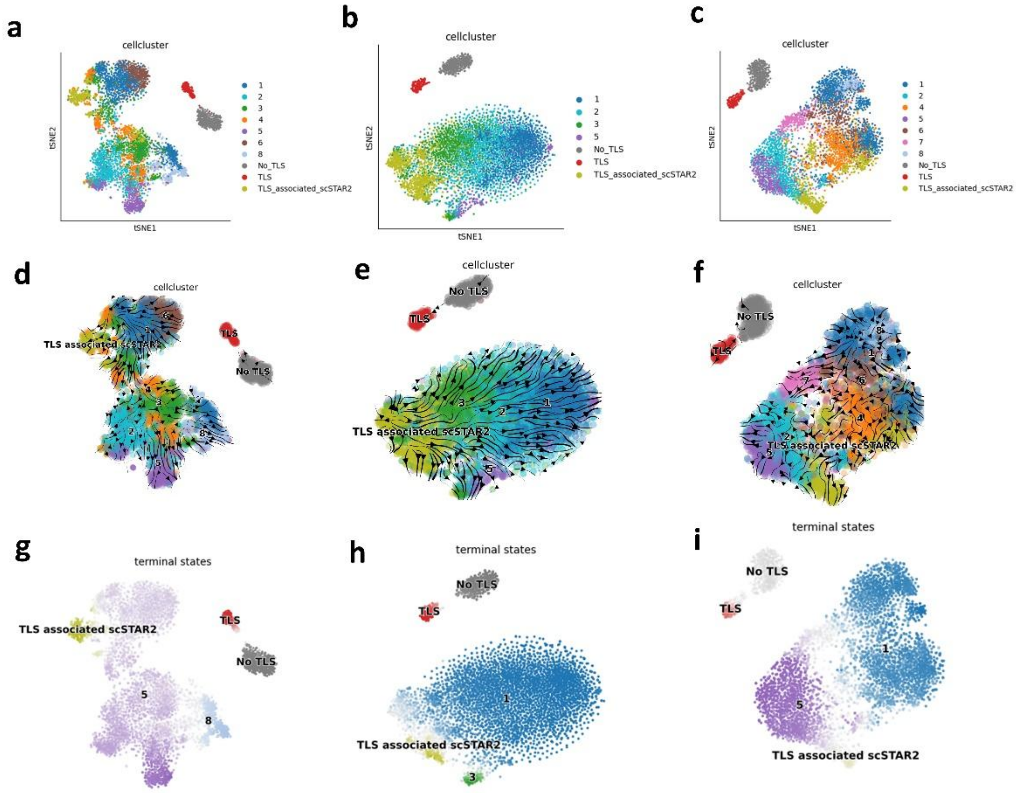


**Figure S5**. The results of cellrank2 applied on original data, slide 1 (a, d, g), slide 2 (b, e, h)– slide 3 (c, f, i).

85

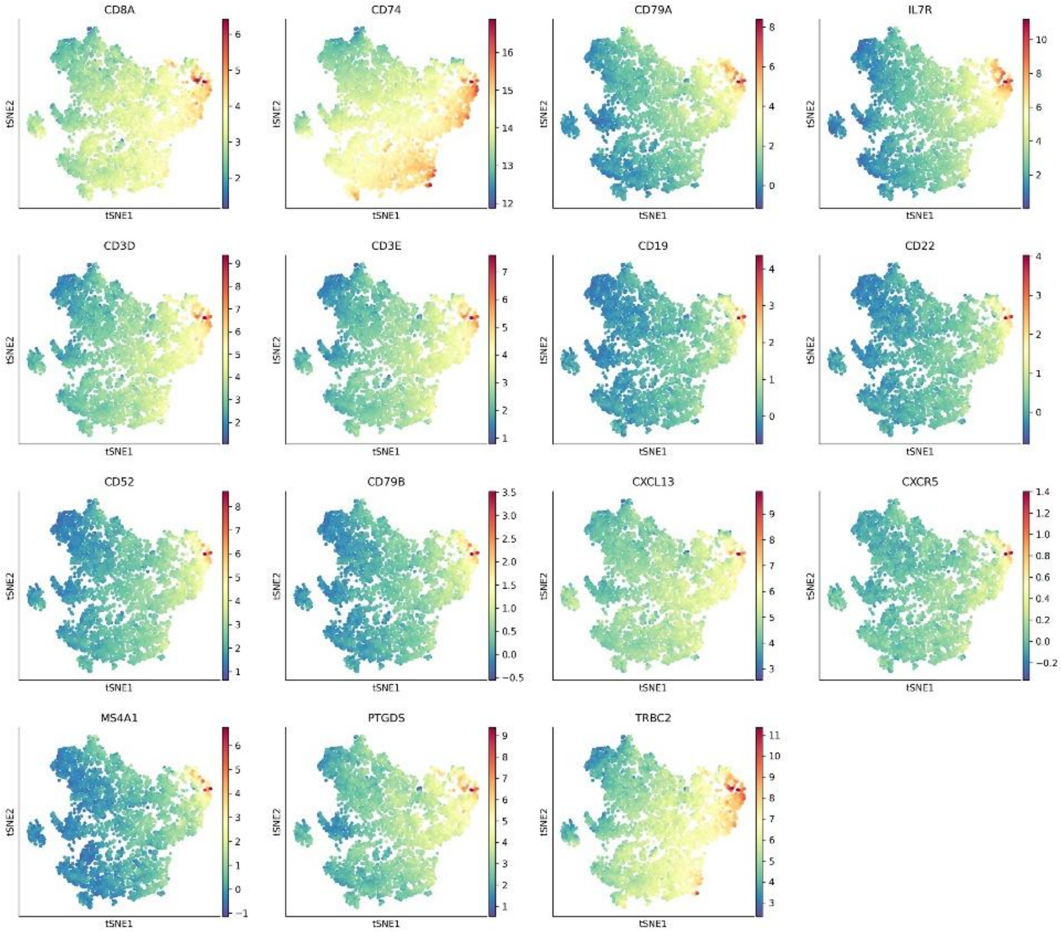


**Figure S6**. The TLS gene expression of each spot of slide1, showed in tSNE map.

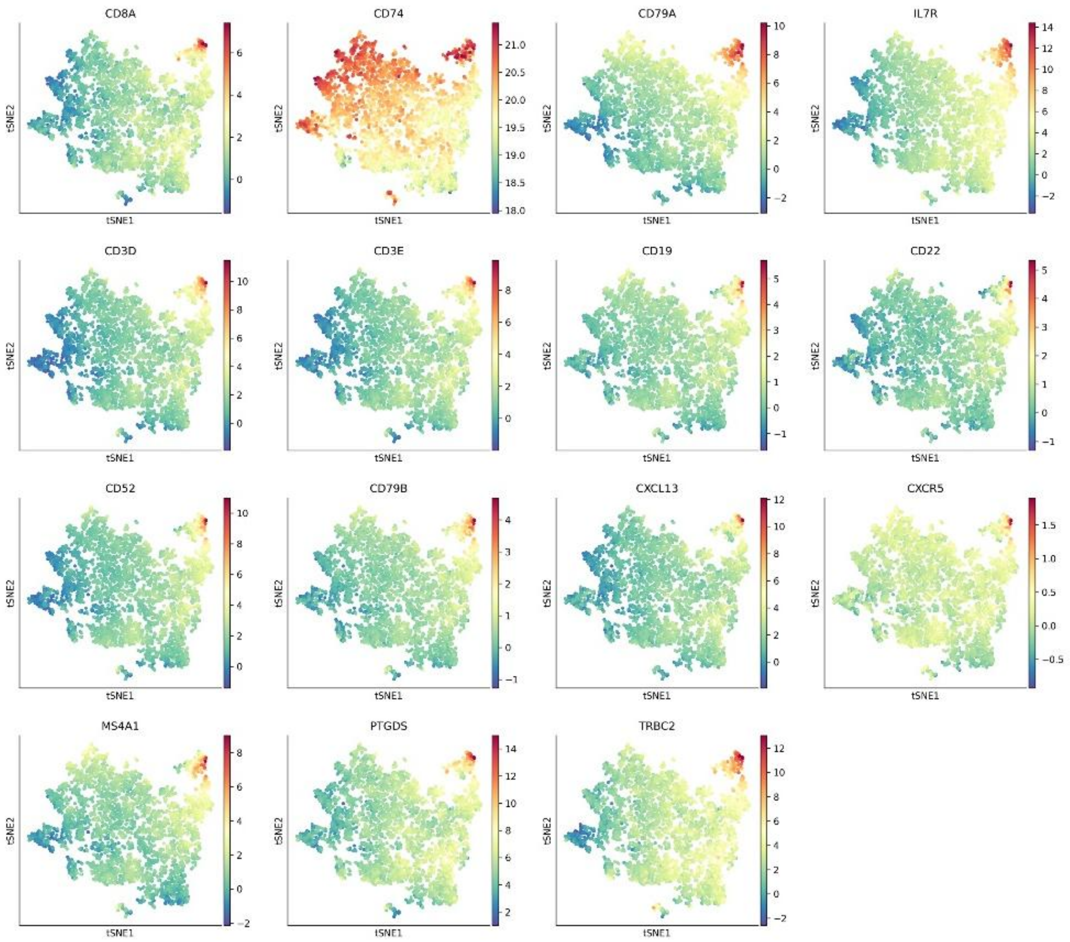


**Figure S7**. The TLS gene expression of each spot of slide2, showed in tSNE map.

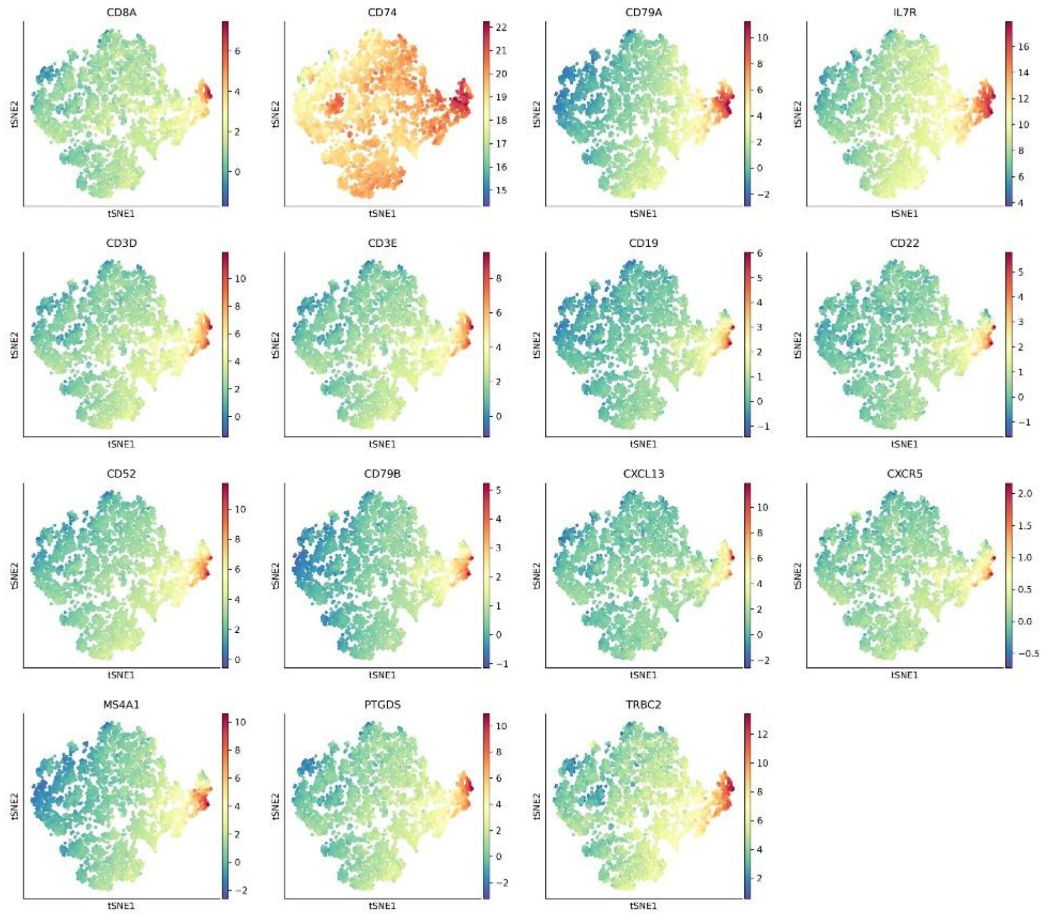


**Figure S8**. The TLS gene expression of each spot of slide2, showed in tSNE map.

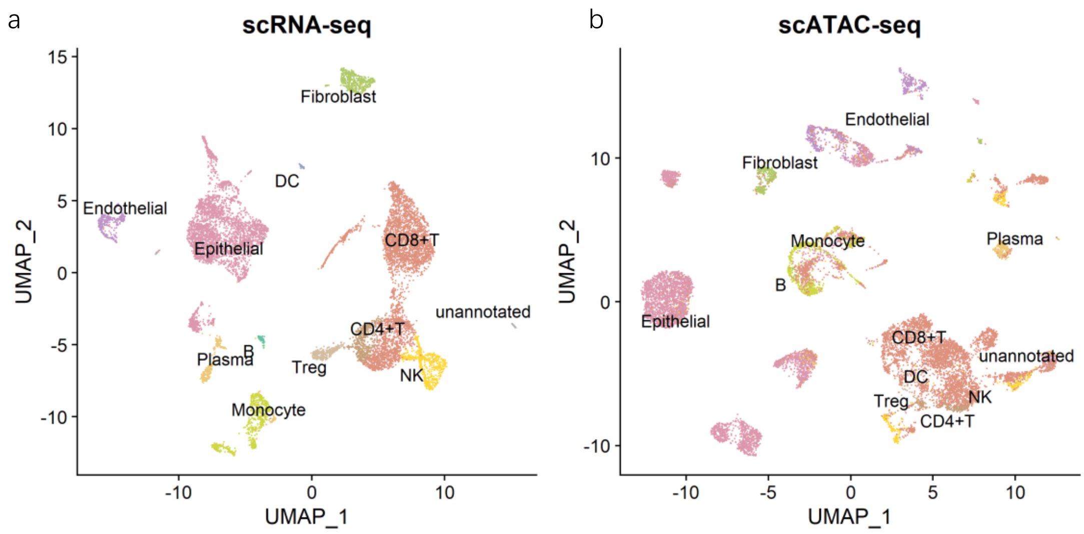


**Figure S9.** ccRCC scATAC-seq and scRNA-seq joint annotation analysis.
